# Adaptive protein evolution through length variation in short tandem repeats

**DOI:** 10.1101/310045

**Authors:** William B. Reinar, Anne Greulich, Ida M. Stø, Jonfinn B. Knutsen, Trond Reitan, Ole K. Tørresen, Sissel Jentoft, Melinka A. Butenko, Kjetill S. Jakobsen

## Abstract

Intrinsically disordered protein regions are of high importance for temperature sensing and immune responses in plants. Tracts of identical amino acids accumulate in these regions and can vary in length over generations due to expansions and retractions of short tandem repeats at the genomic level. However, little attention has been paid to what extent length variation is shaped by natural selection. By environmental association analysis on 2,514 length variable tracts in 770 whole-genome sequenced wild *Arabidopsis thaliana* we show that length variation in glutamine and asparagine amino acid homopolymers, as well as in interaction hotspots, correlate with local bio-climatic habitat. We determined experimentally that the promoter activity of a light-stress gene depended on polyglutamine length variants in a disordered transcription factor. Our results show that length variations impact protein function and are substantially shaped by natural selection. Length variants modulating protein function at a global genomic scale has implications for understanding protein evolution and eco-evolutionary biology.

## Introduction

Hypervariable short tandem repeats (STRs) – also termed microsatellites – are present within gene transcripts and intergenic regions throughout the Tree of Life (*1*). Within the coding part of genes, STRs with a unit size of three encode repeated amino acids (homopolymers) of different lengths without generating codon frameshifts. A substantial percentage of vertebrate and plant proteins (8%-9%) contain homopolymer tracts encoded by STRs (*2*). Unlike the average protein sequence, homopolymer evolution is driven by a different mode of mutation: DNA replication slippage. DNA replication slippage leads to altered homopolymer lengths over generations. It is known that such length variation may cause disease in humans (*3*). However, little is known regarding the effects of benign homopolymer length variation on protein function and structure, and if length variants relate to environmental adaptations.

Interestingly, homopolymer tracts tend to accumulate in protein regions lacking a stable structure, termed intrinsically disordered regions (IDRs) (*4*). In *Arabidopsis thaliana* (henceforth Arabidopsis) the importance of IDRs at the phenotypic level was recently demonstrated in relation to temperature sensing, osmotic stress sensing, and the plant immune response (*5–7*). Furthermore, homopolymer length variation among Arabidopsis genomes has been demonstrated (*8, 9*).

Here, we address if the generation of homopolymer length variants (hypervariability) provides an important, yet unexplored mode of protein evolution in plants and other organisms. We use the 1001 Genomes Consortium dataset (*10*) of whole-genome sequenced wild Arabidopsis to estimate the length of every homopolymer tract in 770 wild accessions covering a wide geographical and environmental spectrum. We combine environmental association analysis with IDR protein structure predictions to investigate if natural selection shape homopolymer length variation and furthermore, experimentally uncover a role for length variable polyglutamine (poly-Q) tracts in light-stress tolerance.

## Results

### Coding STRs predominantly encode structurally disordered homopolymers

We analyzed all protein coding DNA sequences in the Arabidopsis TAIR10 reference genome and found the most common STR-encoded homopolymers to be polyglutamates (poly-E), polyserines (poly-S) and polyaspartates (poly-D) (**Figure 1A, S. Table 1**). To address the hypothesis that length variation in STRs could tune intermolecular interactions by introducing structural changes in disordered protein stretches, we assessed if homopolymers in Arabidopsis proteins coincided with predicted IDRs, as well as with predicted disordered protein-protein, protein-DNA and protein-RNA interaction regions (PPIs, DPIs and RPIs, respectively). The analysis scheme is depicted in **Figure 1B**. To assess the extent of over-or underrepresentation of the homopolymers in IDRs and/or in disordered interaction regions we used two-sided Fisher’s exact tests. The results are shown in **Figure 1C** and **S. Table 2**. We found that 11 out of 20 different amino acid homopolymer tracts to be overrepresented in IDRs, broadly mirroring patterns identified in mammalian and avian proteins (*4*). In PPI regions predicted by DisoRDPbind (see legend **Figure 1**) the strongest enrichments were for poly-E (odds ratio: 9.0), poly-Q (odds ratio: 8.7), and poly-D tracts (odds ratio: 7.6). This differed from the strongest enrichments in MoRFpred (see legend Fig. 1) PPI regions (polymethionine, odds ratio: 9.1), and ANCHOR2 (see legend Fig. 1) PPI sites (polyhistidines, odds ratio: 6.1). In DPI regions, polylysine (poly-K) tracts were strongly enriched (odds ratio: 21.0) and in RPI regions, the strongest enrichment was for polyarginine tracts (odds ratio: 6.1). Importantly, these results show that STR-encoded homopolymers are non-randomly distributed with regard to IDRs and disordered interaction regions. We previously genotyped STRs in the Arabidopsis accessions released by the 1001 Genomes Consortium and defined 770 accessions as a high quality subset (*9*). We used this subset (**S. File 1**) to further explore the characteristics of length variable coding STRs. Notably, we found that polar (S, threonines [T], asparagine [N], and Q) and negatively charged (D, E) homopolymers had a high number of estimated STR length variants present in the population (**Figure 1D**). To better grasp the biological relevance of proteins with length variable homopolymers in IDRs and/or in disordered interaction regions, we performed network and Gene Ontology (GO) analysis of the proteins via the STRING database (*11*). The network and GO analyses showed that the protein set (1,850 proteins) were more biologically connected (including physical interactions, co-mentioning in PubMed abstracts and co-expression) than what was expected in the Arabidopsis background (5,995 edges in the network compared to 4,252 expected edges, *P* value < 1.0e^-16^) (**S. File 2**, **S. Table 3**). The top enriched biological process GO terms included heterochronic regulation of development (6 proteins), circadian rhythm (20 proteins), regulation of salt stress responses (12 proteins) and response to chitin (21 proteins). Notably, we found a high number of TCP family proteins (TCP2, TCP3, TCP4, TCP7, TCP10, TCP13, TCP14, TCP15 and TCP24) in the network (9 of the 24 TCP members, Protein Families [PFAM] enrichment *P* value: 0.01, **S. Table 3**). TCP proteins are known to contain IDRs and to be responsive to environmental stimuli (*12, 13*). Next, based on gene expression analyses from previous work (*9*), we investigated if length variable STRs encoding homopolymers in IDRs or disordered interaction regions were more frequently associated with the expression of the gene it resides in compared to other coding STRs. Supporting that DNA-binding homopolymer tracts could function as autoregulatory gene expression modulators, we found that homopolymers likely to bind DNA (i.e., in DPI regions) were significantly overrepresented (**Figure 1E, S. Table 4**). Furthermore, the tools predicted elevated protein-protein binding propensity in the poly-Q tract of the thermosensor region in EARLY FLOWERING 3 (ELF3), as well as in the poly-E tract of ALFIN-LIKE 6 (AL6), where we previously demonstrated an effect on protein-protein interactions (*9*). This supports that the prediction tools can point to known biologically interesting regions. PPI, DPI and RPI predictions for ELF3, AL6, and other example proteins are shown in **S. Figure 1**.

**Figure 1.**
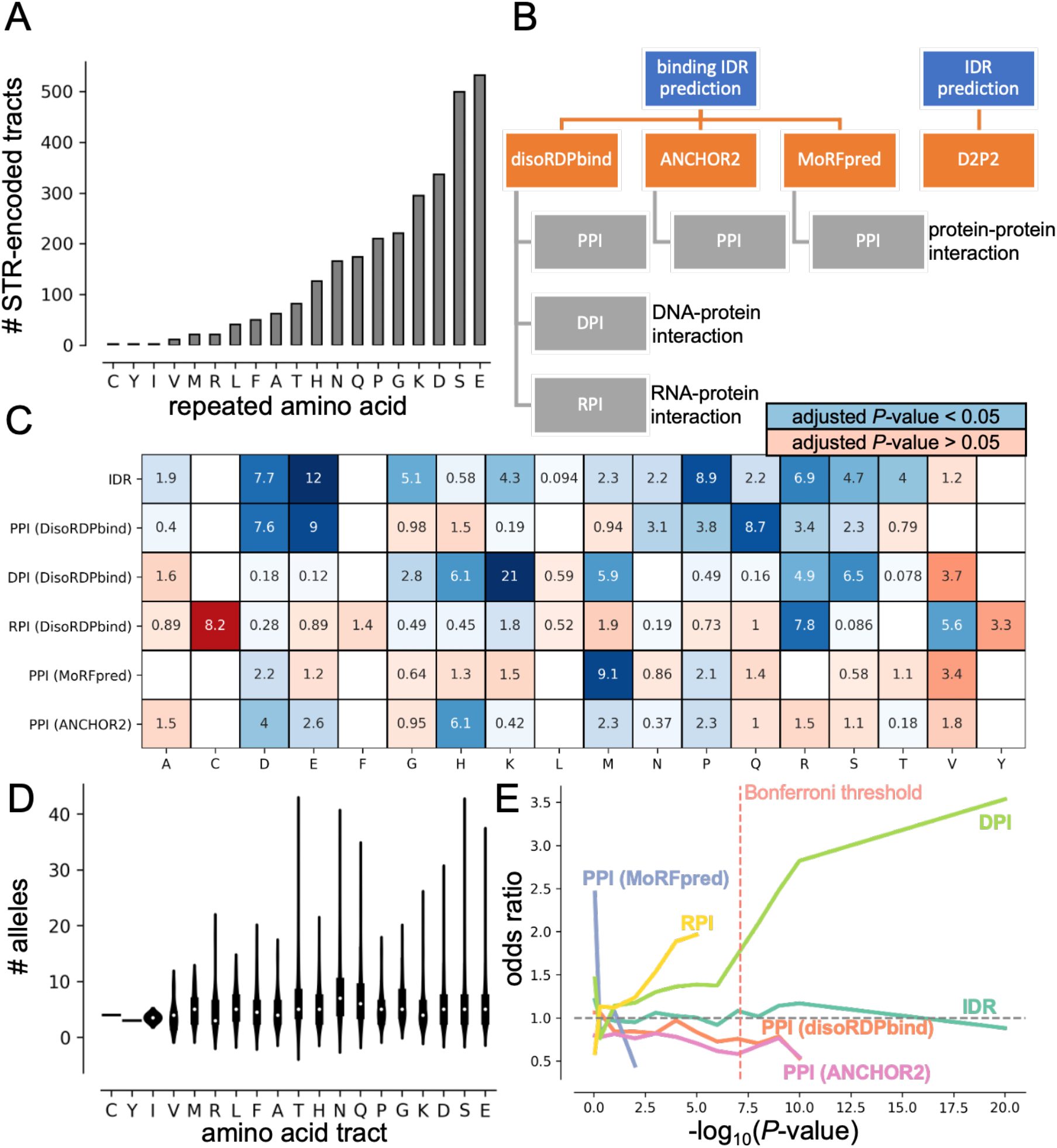
Protein coding short tandem repeats (STRs) often encode disordered regions in *Arabidopsis thaliana* proteins. (**A**) The number of STR-encoded homopolymer tracts per type of amino acid in the *Arabidopsis thaliana* reference (TAIR10) protein sequences. (**B**) Outline of prediction tools employed to define intrinsically disordered regions (IDRs) and disordered interaction regions (protein-protein interaction: PPI, DNA-protein interaction: DPI and RNA-protein interaction: RPI). (**C**) The degree of over-or underrepresentation of homopolymer tracts encoded by reference genome STRs within the predicted (IDR, PPI, DPI and RPI) regions. Box color shading is scaled with odds ratio values and are in blue tones if the Fisher’s exact test *P-*value was significant at the 0.05 level after Bonferroni correction, and in red tones otherwise. (**D**) The violin plots show distributions of the estimated number of STR alleles in 770 different *Arabidopsis thaliana* accessions (S. File 1). (**E**) STR-encoded homopolymers that overlap predicted disordered DNA-binding regions were overrepresented in being associated with autoregulatory gene expression compared to other protein coding STRs (S. Table 4). The red, dashed line indicates the *P-*value threshold after adjusting for multiple tests (Bonferroni) in the gene expression analysis. Amino acid legend: A: alanine C: cysteine, D: aspartate, E: glutamate, F: phenylalanine, G: glycine, H: histidine, K: lysine, M: methionine, N: asparagine, P: proline, Q: glutamine, R: arginine, S: serine, T: threonine, V: valine, Y: tyrosine. Note that no STR-encoded tracts of tryptophane (W) were detected in the TAIR10 reference proteins.

### ELIP1 promoter activation depends on the length of poly-Q tracts in TCP14

Several members of the TCP transcription factor family had STR-encoded homopolymers predicted to be in IDRs and/or in disordered interaction regions (**S. Table 3**, **S. File 2**). One member of this family, TCP14, activates the photoprotective EARLY-LIGHT INDUCED PROTEINS (ELIP*s*), which are involved in plant responses to high light stress *(14–16).* The expression of *ELIP1* is directly regulated by the binding of TCP14 to *up1* elements in the *ELIP* promoter regions. TCP14 acts epistatic to the DnaJ-like zinc finger domain-containing transcriptional regulator ORANGE (OR). OR physically interacts with TCP14 in the nucleus and represses its transactivation activity leading to reduced transcriptional levels of *ELIP1*, reduction in chlorophyll biosynthesis and delays of thylakoid membranes in plastids of germinating cotyledons. This repression decreases upon illumination when the nuclear localization of OR is diminished and accumulation of TCP14 in the nucleus derepresses chloroplast biogenesis during hypocotyl de-etiolation by increased *ELIP1* transcription (*15*). We found TCP14 to contain poly-Q tracts that overlapped predicted disordered PPI regions (**S. Figure 1**) and that varied in length in different Arabidopsis accessions. We experimentally tested the functional relevance of poly-Q tracts in TCP14 by quantifying the ability of the different homopolymer length variants to activate transcription of the *ELIP1* gene. The two poly-Q tracts we focused on overlap with predicted disordered PPI sites (**Figure 2A**), and we envisioned that length variation in these poly-Q tracts would influence the ability of TCP14 to activate *ELIP1.* We tested our hypothesis using *TCP14* variants from accessions that only differed in these two tracts in the TCP14 protein sequence. The Col-0 accession had seven repeated Qs in the first tract and three repeated Qs in the second tract (TCP14-7Q-3Q). The Bulgarian accession CS77239 had four repeated Qs in the first tract and six repeated Qs in the second tract (TCP14-4Q-6Q). No other amino acid level differences were present between the two accessions (**S. File 3**). To isolate the total effects of the first and the second tract, we synthesized and experimentally tested all the remaining combinations (7Q-6Q and 4Q-3Q). To verify that the activity of the *ELIP1* promoter *(pELIP1)* was significantly altered by the length of the TCP14 poly-Q tracts we performed a luciferase assay in *Nicotiana benthamiana (N. benthamiana)* leaves where the luciferase luminescence signal (relative light units [RLU]) was a determinate of promoter activity. We co-infiltrated an *ELIP1p:LUC* construct together with the different TCP14 proteins coupled to green fluorescent protein (GFP) in *N. benthamiana* leaves (TCP14-Qn-Qn). An estradiol-inducible 35S promoter was used to drive expression of the TCP14-Qn-Qn-GFP variants to achieve comparable expression levels for the fusion proteins. TCP14-7Q-3Q-GFP, TCP14-4Q-6Q-GFP, TCP14-7Q-6Q-GFP and TCP1-4Q-3Q-GFP localized to the nucleus in *N. benthamiana* leaf cells (**Figure 2B**). As negative controls, we measured the luciferase luminescence signals in assays containing the constructs without adding estradiol, as well as assays without *ELIP1p:LUC* (Figure 2C). Our results show that a simultaneous length change in both tracts produced significantly different luciferase outputs (7Q-6Q vs. 4Q-3Q: Welsch T-test *P* value: 0.007 and 4Q-6Q vs. 7Q-3Q: Welsch T-test *P* value: 0.03), and that the last tract yielded a significant difference only when the first tract was 7Q (7Q-6Q vs. 7Q-3Q: Welsch T-test *P* value: 0.0008). Changes in only the first tract did not produce any significant differences (Figure 2C). We conclude that both tracts are involved with *ELIP1* promoter activation, but that the last tract is more important. These results show that natural allelic variations in the poly-Q tracts alters *ELIP1* promoter activation, and given the important role of ELIP1 in light stress toleration, this most likely reflects the ability of different accessions to respond to light stress.

**Figure 2.**
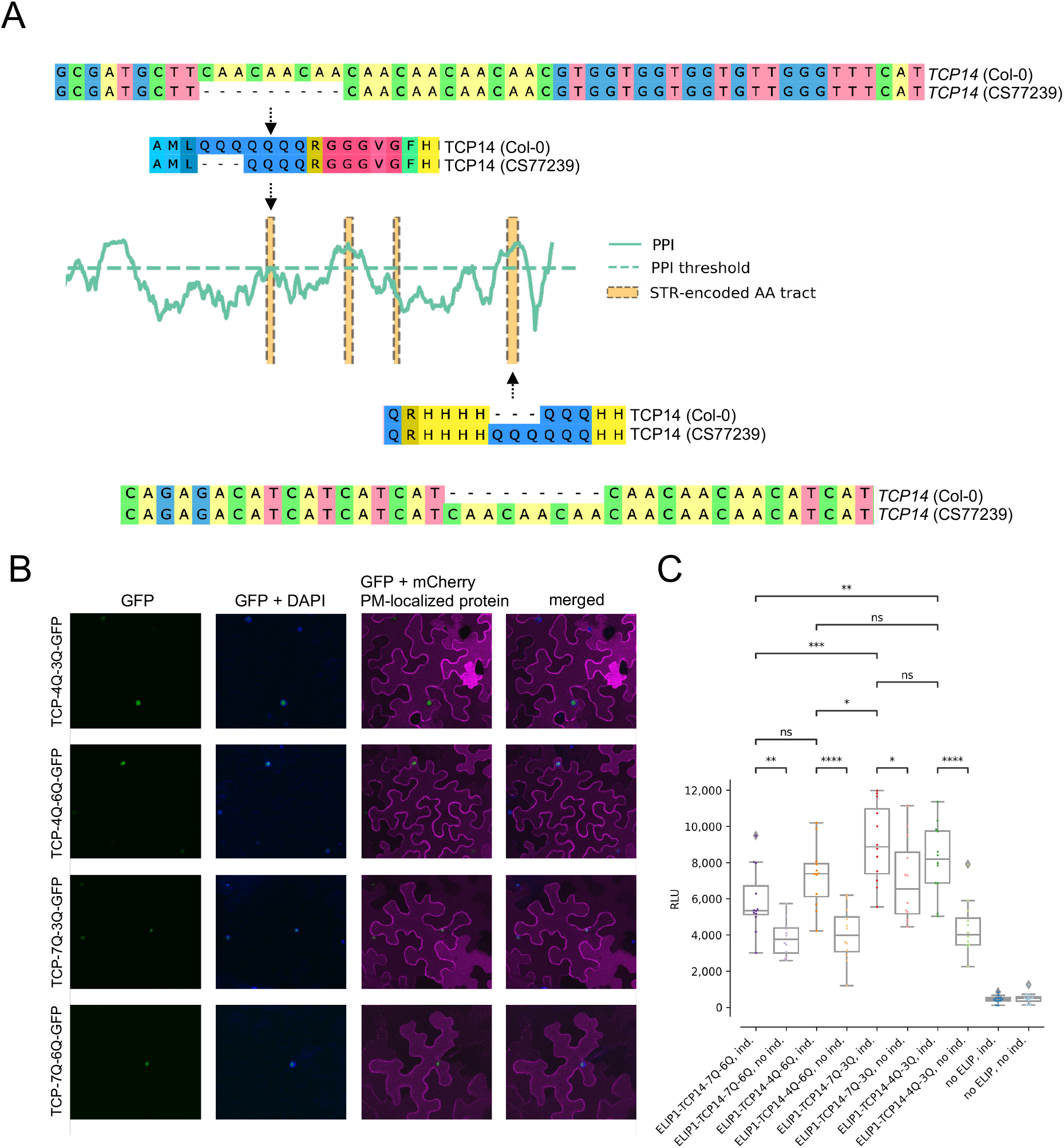
Natural allelic variation in two short tandem repeat (STR)-encoded polyglutamine (poly-Q) tracts in TCP14 influences the activation of the *ELIP1* promoter. **(A)** The alignments show excerpts of the *TCP14* (Col-0) and *TCP14* (CS77239) gene and protein sequence, highlighting the two poly-Q tracts with natural allelic length variation. The cartoon shows the location of the poly-Q tracts relative to the predicted protein-protein binding propensity of the TCP14. **(B)** Transient expression of synthesized TCP14 (TCP14-4Q-3Q-GFP), TCP14 from accession CS77239 (TCP14-4Q-6Q-GFP), TCP14 from Col-0 (TCP14-7Q-3Q-GFP), and synthesized TCP14 (TCP14-7Q-6Q_GFP) in *Nicotiana benthamiana* leaves. The images show that TCP14-4Q-3Q-GFP, TCP14-4Q-6Q-GFP, TCP14-7Q-3Q-GFP, and TCP14-7Q-6Q-GFP localize to the nuclei. A protein known to localize to the plasma membrane (PM) was used to outline the PM of the cells. **(C)** The luciferase output (relative light units [RLU]) of the *ELIP1* promoter is dependent on the Q tracts in TCP14. The differences in luciferase output show that all TCP14 variants activate the *ELIP1* promoter, and that when both tracts are changed, so is the strength of the activation. The asterisks indicate the result of Welch T-test (****: *P* value < 0.0001, ***: *P* value < 0.001, **: *P* value < 0.01, *: *P* value < 0.05, ns: *P* value > 0.05). Ind.: induction.

### Homopolymer tract lengths vary along temperature-related gradients

Next, we explored the relationship between natural allelic variation in protein coding STRs and the environmental origin of the accessions. To perform such analyses, we retrieved environmental measurements gathered by Ferrero-Serrano and Assmann (2019) that covered the 770 accessions(*17*). 88 numerical variables had complete measurements for all the 770 accessions (**S. File 4)**. As many of these variables were highly correlated, we decomposed the information to seven principal components (PCs) that together explained 80% of the total variation (**S. Figure 2)**. The top correlations between the individual environmental variables and PC axes 1-7 are shown in **S. Table 5**, and the accessions’ values along the axes are available in **S. Table 6** (PCA plot shown in **S. Figure 3**). We regressed the seven environmental PC axes on our estimates of allelic variation in 2,514 protein coding STRs, one site and one environmental axis at a time (**S. File 5**). The accessions’ original sampling location colored by their position along PC1 and PC2 are shown in **Figure 3A.** As a control, we used mock STR genotypes (**S. File 6**). We separately treated STR alleles as continuous or categorical values to assess if linear or non-linear effects best explained the environmental variation. In 24% of tests with multiallelic STRs, non-linear effects better explained the environmental variation compared to linear effects. For further analysis, we kept the top associations (by *P*-value) per PC axis and per STR. Of the 2,514 protein coding STRs, 948 had length variation best associated with PC axis 1 (**Figure 3B, S. File 5**). This axis mostly represent variation in spring maximum temperatures (**S. Table 5**). At the highest confidence levels (i.e. lowest *P*-values), associations with PC axis 2 were more common (**Figure 3B, S. File 5**). PC axis 2 best represents variation in net primary production and temperature seasonality (**S. Table 5**). None of the mock genotypes produced significant associations with any PC axes after multiple test (Bonferroni) correction (**S. File 6**). With regard to specific homopolymer tract types, poly-N, poly-Q, and poly-E tracts were overrepresented in producing higher confidence associations with PC axis 1, and poly-E and poly-S tracts were driving the higher confidence associations with PC axis 2 (**Figure 3C**). Notably, the top candidate proteins containing poly-N, poly-Q, poly-E or poly-S tracts with environmentally associated length variation included proteins known to be involved in regulating responses to abiotic and biotic stress (**S. Figure 4**). Note that we cannot completely rule out that the associations are confounded with the population structuring along the environmental gradients (**S. Figure 5**). However, when controlling for the pairwise genetic similarity with putative neutral markers (**S. Figure 6**), *P*-value distributions were right-skewed (**S. Figure 7**), and 23 coding STRs were still significant after Bonferroni correction (**S. File 7**).

**Figure 3.**
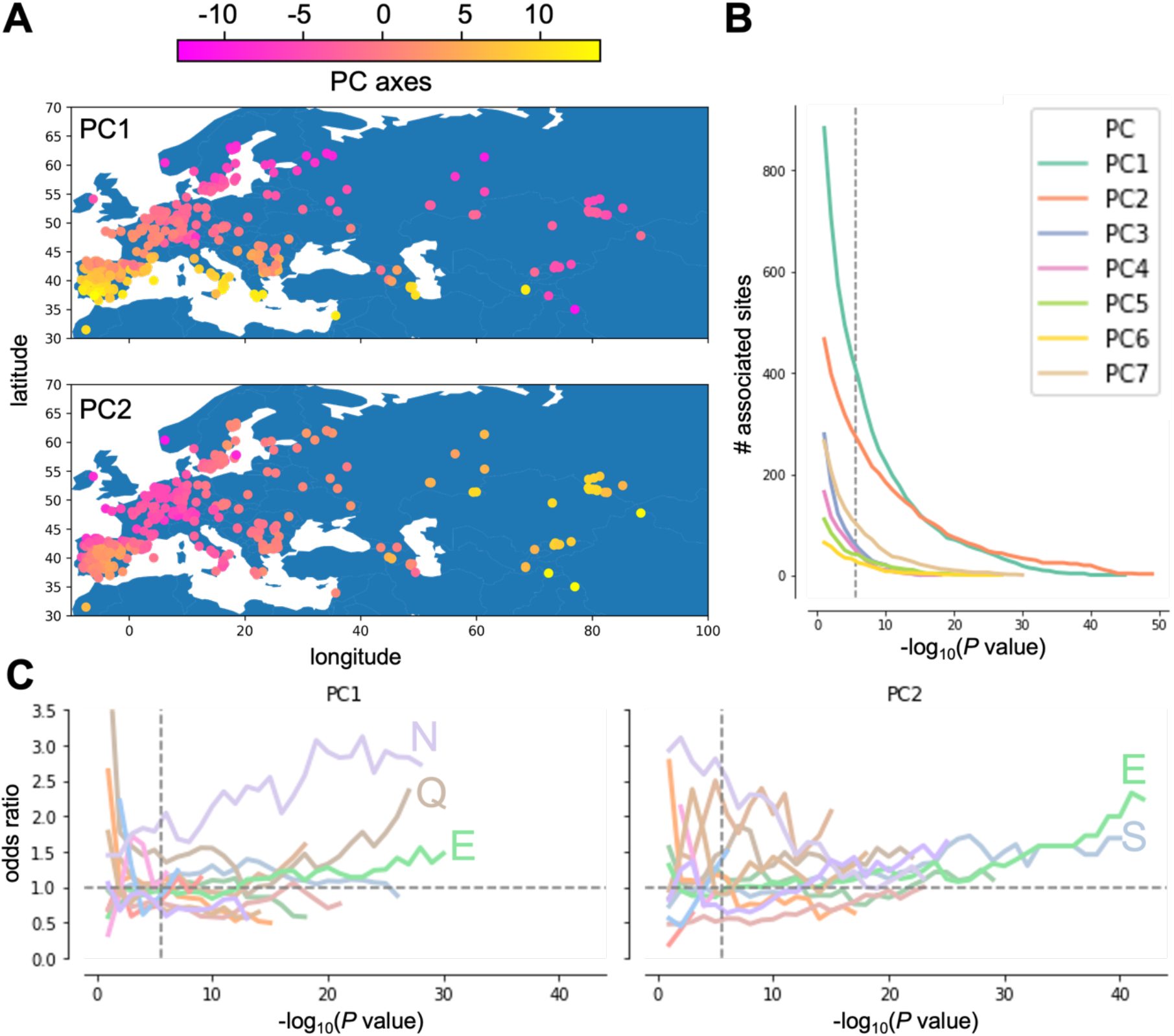
Natural allelic length variation in short tandem repeats (STRs) encoding polyasparagine (poly-N), polyglutamine (poly-Q), polyserine (poly-S) and polyglutamate (poly-E) tracts associate with environmental gradients related to temperature and productivity. (**A**) The maps show the sampling origins of the 770 *Arabidopsis thaliana* accessions. Samples are colored by their position on the environmental principal component (PC) axes 1 and 2 (see colorbar). (**B**) The number of protein coding STR sites (y-axis) associated with the environmental PC axes 1-7 as a function of the confidence of the environmental association (-log_10_ *P* values). (**C**) The line plots show the odds ratios (y-axis) resulting from one-sided Fisher’s exact test of dependence between the type of amino acid and its association with PC axis 1 (left panel) and PC axis 2 (right panel). The x-axis show the confidence of the association (-log_10_ *P* value) with the PC axis. The odds ratios of association with the other PC axes are shown in **S. Table 7**.

### Environmental correlations partly driven by disorder and binding propensities

Next, we investigated if length variation in disordered homopolymer tracts is prone to natural selection. For every homopolymer tract we treated the squared correlation (R^2^) between environmental variation (PC axes 1–7) and the homopolymer length variation as a naïve estimate of natural selection, *S* (**S. File 5**). In a multiple regression model, we tested if the homopolymer tract type (poly-N, poly-S, etc.) significantly contributed to explain variation in *S* (**Figure 4**). In the same model, we included presence/absence of homopolymers in IDRs and/or IDR interaction regions (PPIs, DPIs, and RPIs) as explanatory variables (**S. File 8**). Combined, these variables explained 1.2% of the variation in *S* (**S. Table 8**). At the multiple test adjusted significance threshold (α: 2.8 × 10^-6^), poly-N tracts significantly contributed to elevated *S*-values (T: 5.9, *P*-value: 4.2 × 10^-9^). At this threshold, the presence of homopolymer tracts in IDRs and in DPIs were significant factors as well (IDR: T: 3.2, *P*-value: 6.8 × 10^-4^; DPI: T: 3.3, *P*-value: 5.0 × 10^-4^). At a less strict significance threshold (α: 0.05), poly-Q and presence in PPIs contributed to elevated *S-* values (poly-Q: T: 3.0, *P*-value: 0.003; PPI: T: 2.6, *P*-value: 0.009), and poly-K tracts notably contributed to a decrease in *S*-values (T:-2.5, *P*-value: 0.01). We repeated the test with log_10-_transformed R^2^-values and got similar results (**S. Table 8**). Although the simple models used here only explains a small fraction of variation in *S* our results clearly indicate that disordered homopolymer tracts, as well as disordered homopolymer tracts with predicted DNA and prote-inbinding properties are likely targets of natural selection.

**Figure 4.:**
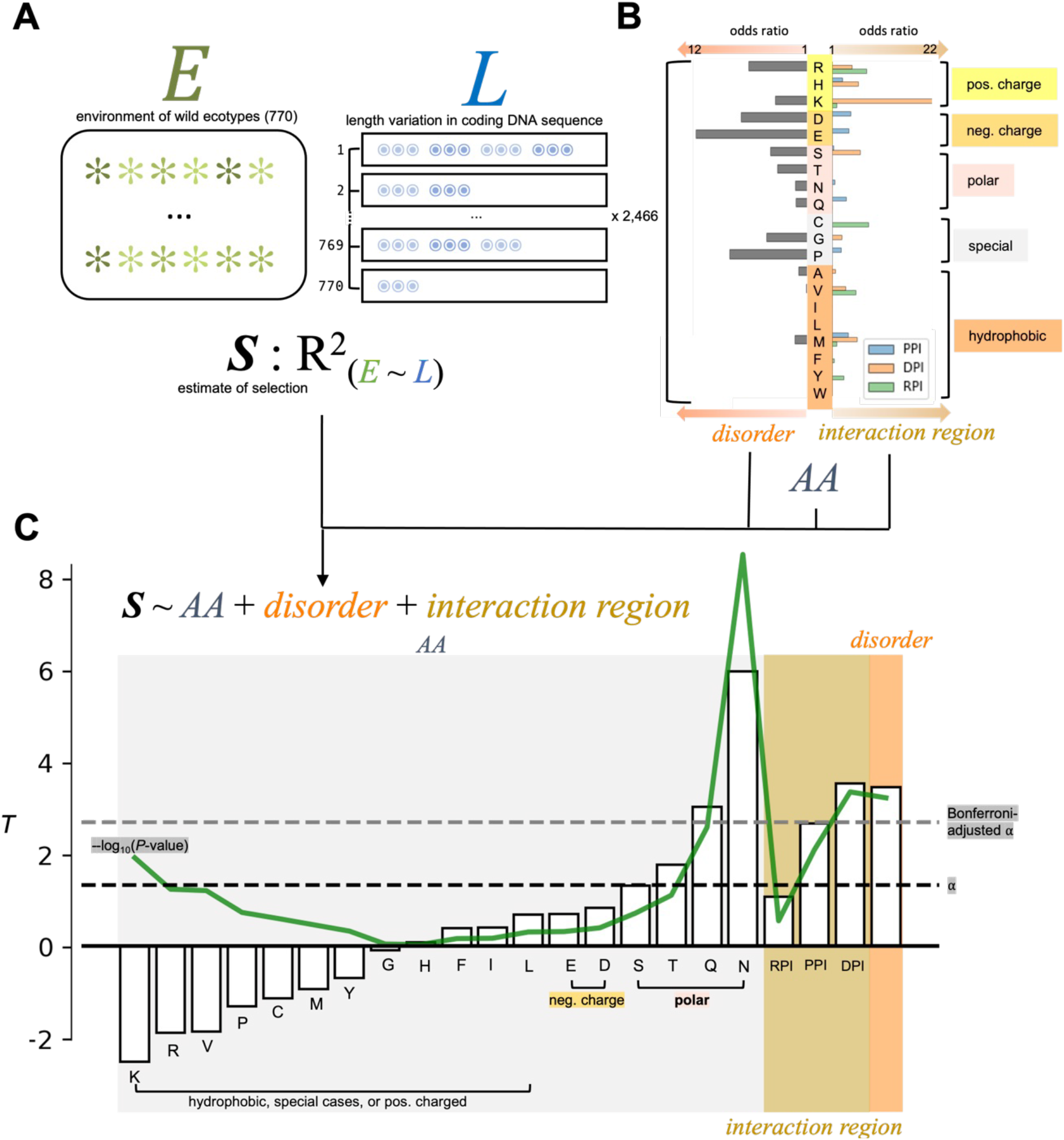
Length variation in disordered homopolymer tracts linked to variation in environmental gradients. (**A**) As an estimate of selection on homopolymer tracts, S, we used the squared correlation (R^2^) between homopolymer tract length variation and environmental gradients (environmental PC axes 1-7) (S. File 5). *‘* served as the response variable in a multtple regression model. (**B**) Homopolymcrs differs in what amino acid is repeated *(AA),* and in the extent that traces are found in intrinsically disordered regions (disorder) and in disordered regions predicted to partake in protein-nucleic acid interactions (DNA or RNA) or protein-protein interactions. These variables served as explanatory variables in the model. Note that the effect of*AA*was relative to the baseline *AA,* set to be alanine. (**C**) The bar chart shows die ‘*/*-statistic (y-axis) of each term (x-axis) in the multiple linear regres sion moTel. The green line shows the-log_10_(*P*-value). The black, dashed line denote the chosen significance level, α, at 0.05. The gray, dashed line denotes the Bonferroni-adjvsted *α* (2.8 × 10^-6^). Amino acid legend: K: lysine, R: arginine, V: valine, P: proline, C: cysteine, M: methionine, Y: tyro sine, G: glycine, H: histidine, F: phenylalanine, I: isoleucine, L: leucine, E: glutamate, D: aspartate, S: serine, T: threonine, Q: glutamine, N: asparagine.

## Discussion

In this study we performed a global analysis of protein coding STRs in wild Arabidopsis accessions to explore how coding length variation have the potential to affect protein function, and how length variations relate to environmental conditions.

Our protein structure analyses indicated that coding STRs promote disorder in protein structure, and that STRs in disordered regions often are predicted to bind other proteins, DNA or RNA (**Figure 1C**). The non-random distribution of STR-encoded amino acids with regard to protein structure implies that protein level length variation, caused by random replication slippage on the DNA level, predominantly affects regions of proteins predicted to be disordered. By utilizing the wealth of knowledge resulting from decades of functional studies with Arabidopsis as a model plant, our protein network enrichment analysis (**S. Table 3**) revealed that proteins encoded in part by structural disorder-promoting STRs are not a random subset of the Arabidopsis proteome, but heavily involved in circadian rhythms, defense responses and responses to salt stress, all crucial features for adaptation to microhabitats. The results of our experimental approaches on TCP14, support that the prediction tools pointed to functional disordered amino acid tracts within the TCP14 protein sequence (**Figure 2**). The Q tracts in TCP14 were predicted to have elevated protein-protein binding propensities, making it likely that differential downstream interacting *ELIP1* promoter activation was caused by an altered protein-protein interaction between TCP14 and OR, as OR is known to bind TPC14 and represses its ability to activate the *ELIP1* promoter(*15*). Further experiments on TCP14 would be necessary to fully test this hypothesis, and follow-up experiments on additional candidate tracts, similar to Jung et. al (2020), will be key to understanding the extent of the phenotypic consequences of protein coding STR length variation.

Here, we have demonstrated that length variation in coding STRs is associated with various environmental adaptations in wild Arabidopsis accessions. Based on the knowledge from other plant, fungal, animal and bacterial species, coding STRs are found at high frequencies throughout the Tree of Life (*1, 2,18–24*)—and there are some published examples of functional consequences of tract length variation on specific proteins. These examples include a variable glycine-threonine tract that maintains the circadian rhythm in response to temperature in *Drosophila* (*25*). In yeast, a variable poly-Q tract in a transcriptional regulator protein influenced fitness under different growth regimes (*19*). A variable poly-Q tract in combination with a variable poly-A tract facilitate swift evolution of limb and skull morphology in canids *(26).* Arabidopsis is however, one of the few species where a large collection of whole-genome sequenced wild specimens (adapted to very different local environments) is available. This made it possible to perform a global analysis of homopolymer tract length variation and to find new candidate proteins exhibiting environmentally associated length variants. In an even further and broader perspective, the nature of STR length variation somewhat resembles that of epigenetic changes (epimutations), in that both STR length variations and epimutations occur at a rapid mode, and both STR variation and epimutations may be involved in local adaptation. However, the transgenerational effect of epigenetics is not completely clear, while STR mutations acting at the DNA level are inheritably stable. In turn, this is likely to provide rapid adaptation under specific (and even recurrent changing) selection regimes due to the high length mutation rates and the standing variation in the populations.

Our environmental association analyses showed that specific homopolymer tracts (particularly poly-Q, poly-N, poly-E and poly-S), as well as disordered tracts predicted to bind other proteins and DNA had length variation strongly associated with complex environmental gradients (**Figure 3C, Figure 4**), and that many of the candidate proteins are known to be involved in responses to biotic and abiotic stress, including SEUSS-LIKE 2, MYB DOMAIN PROTEIN 68, ENHANCER OF VASCULAR WILT RESISTANCE and SCREAM2 (**S. Figure 4**). As such, our data strongly suggest that natural selection has driven the STR length variation between ecologically adapted populations and consequently the allelic variation in these tracts as a response to differences in temperature, temperature variability and other more complex gradients. Furthermore, the presence of such tracts in proteins facilitates a rapid fine-tuning of protein function in plants, and likely other groups of organisms since coding STRs are ubiquitous throughout the Tree of Life.

## Materials and Methods

### Disorder and disordered binding site predictions

We downloaded *A. thaliana* reference (TAIR10) proteome disorder consensus predictions from d2p2 (*27*). d2p2 itself uses nine different predictors: VLXT, VSL2b, PrDOS, PV2, IUPred-S, IUPred-L, Espritz-N, Espritz-X and Espritz-D *(28–33).* d2p2 requires that 75% of tools should agree in order to designate disorder. From these consensus ranges, we generated Browser Extensible Data (BED) files. To find overlaps with STR-encoded amino acids, we first ran Tandem Repeats Finder with standard options (*34*) on the TAIR10 coding DNA sequences retrieved from arabidopsis.org. This allowed us to map the STR on the coding DNA transcript level to the protein sequence. As such, we generated another BED file containing amino acids encoded by STRs. We used Pybedtools with the ‘intersect’ command to assess overlap of STR-encoded amino acids and the predicted regions. Pybedtools is a Python wrapper of BEDTools (*35, 36*). For predicting disordered binding regions, we used DisoRDPbind predictions, MoRFpred predictions and ANCHOR2 predictions (*37–39*). For retrieving DisoRDPbind predictions, we accessed the DisoRDPbind web server (http://biomine.cs.vcu.edu/servers/DisoRDPbind/). To retrieve MoRFpred predictions, we uploaded protein sequences to the fMoRFpred web server available at http://biomine.cs.vcu.edu/servers/fMoRFpred/ (*40*). To retrieve ANCHOR2 predictions, we ran the IUApred2A tool locally (*41*). The different tools’ output result files were parsed to generate BED files, and overlap was quantified using Pybedtools. To link the STR-encoded homopolymer coordinates back to the genomic coordinates of the STR, we matched transcript IDs from the coding DNA sequence FASTA file to the genome annotation GFF file (from arabidopsis.org) and then located the identical motifs detected by Tandem Repeats Finder in the TAIR10 genome sequence and in the coding DNA sequence FASTA file.

### Allelic variation in STRs and neutral SNPs

We previously scored the number of units in each (diploid) STR allele across the Arabidopsis accessions(*9*). An STR-specific variant caller, Haplotype inference and phasing for Short Tandem Repeats (*42*) was used to call the variants. We used the same STR unit counts in this study, but for an expanded set of accessions (770), and only STRs that encoded homopolymer tracts. We constructed the putative neutral SNP matrix for the 770 accessions as in our previous study (*9*). In brief, we drew random, common (major allele frequency < 0.9) intergenic SNPs that we pruned for linkage disequilibrium using the ‘locate_unlinked’ function of the Python package scikit.allel (parameters: size = 1000, step = 20, threshold = 0.1)(*43*). From these SNPs, we calculated the pairwise correlation between accessions and standardized the output matrix.

### Bioclimatic variables

We retrieved the full set of environmental variables retrieved by Ferrero-Serrano and Assmann (2019) (*17*) and dropped measurements that were not fully covered in all samples, which left 88 different environmental variables (S. File 3). First, we scaled the variables using the ‘sklearn.preprocessing.scale’ and then the ‘sklearn.decomposition.pca’ functions of the ‘sklearn’ Python module (*44*) to perform the decomposition of the 88 variables.

### Cloning and transient expression of proteins

Arabidopsis accession CS77239 was ordered from Nottingham Arabidopsis Stock Center. Clonal genes of TCP14-7Q-6Q and TCP14-4Q-3Q were ordered from Twist Bioscience. The DNA sequences encoding *TCP14* from Col-0 and *TCP14* from accession CS77239 were cloned in frame with an expression vector containing an 35S estradiol-inducible promoter and a C-terminal fluorescent molecule of GFP (*45*) using the Invitrogen Gateway cloning system. The *ELIP1* promoter (pELIP1) was defined as the region 2,000-bp upstream of the *ELIP1* start codon and amplified from Col-0 genomic DNA. *pELIP1* was cloned into the R4pGWB635 vector *(46, 47)* containing the LUC gene, creating the pELIP1:LUC construct. Primers are listed in S. Table 7. Plasmids were transformed into *Agrobacterium tumefaciens* C58 and further used for transient expression in *Nicotiana benthamiana* leaves following a previously described protocol (*48*).

### Luciferase assay

*N. benthamiana* leaves were co-infiltrated with *ELIP1:LUC* and one of the TCP14 constructs individually. The empty vector pab117 served as a no *ELIP1:LUC* negative control. Leaves were cut into disks of 3 mm in diameter and incubated in a 10-mM estradiol solution for three hours to induce gene expression of TCP14. Uninduced leaf disks for each construct were used as a negative control for TCP14 activation of *ELIP1:LUC.* Leaf disks were individually transferred to wells in a 96-well plate treated with 0,5 mg/ml D-luciferin and kept in darkness for 105 minutes before luminescence was detected using a BioTek Synergy H1 microplate reader (Agilent). For imaging, leave discs were incubated in a 10-mM estradiol and 2,5 μg/ml DAPI solution mix overnight. Images were captured using a Olympus FV1000 Inverted microscope with a Uplan-Apochromat 20x/0.7 WD=0.65 objective.

### Statistical analysis

Fisher’s exact test can be used to test for dependence by calculating how extreme observed differences in ratios are, given no dependence. We used this test with different kinds of counts in this study. In relation to disorder predictions, we used Fisher’s exact test to investigate if STRs and the predicted features were dependent. Given no dependence between two features (for instance between STRs and predicted IDRs), we would not expect a large difference between ratios. As such, we constructed contingency tables and used the ‘fisher_exact’ function of the Python module ‘statsmodels’ to calculate odds ratios and *P* values from contingency tables (*49*). The odds ratio would be 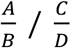 for a given type of amino acid, where A is the count of STR-encoded amino acids in the predicted region, B is the count of amino acids not encoded by STRs in the predicted region, C is the count of STR-encoded amino acids not in the predicted region, and D is the count of amino acids not encoded by STRs and not in the predicted region (see legend **S. Table 2**). We used two-sided Fisher’s exact test, as we were interested in both positive and negative dependence. For investigating dependence of the nature of the repeated amino acid (for instance poly-Q tracts) and associations with the environment we used the same method (see legend **S. Table 7**). Here, we were only interested in positive dependence and performed one-sided Fisher’s exact test (with the option ‘greater’ in the ‘fisher_exact’ function). We also used one-sided Fisher’s exact tests to address if there was dependence between IDRs or predicted IDR binding sites and associations with gene expression (see legend S. Table 4).

We used ordinary linear square regression to test if the environment of Arabidopsis accessions could be predicted by allelic variation in coding STRs. The modeling was performed using the ‘ols’ function of the ‘statsmodels’ Python package. We ran ordinary least square (OLS) regression with the seven PC axes as a response, separately, and each STR separately. In addition, STRs were modeled as categorical variables, which tests for non-linear STR effects. For multiallelic STRs (more than two variants), we compared the non-linear STR effect model with the linear model using the ‘anova_lm’ function of ‘statsmodels’ and kept the *P-*value and R^2^ of the best model. Mock STRs were generated by shuffling the real STR genotypes randomly among the accessions, per STR site, and were modelled in the exact same manner as the real STRs. For purposes of correcting for population structure we used linear mixed models as described in (*9*). Briefly, we used the ‘qtl.scan’ function of the Python package ‘limix’ (https://github.com/limix/limix), setting G as the STR genotypes, Y as the environmental PC axes and K as the standardized genetic correlation matrix. Models with and without G were then compared, and *P-*values were calculated by comparing the likelihood ratios of the two models.

We first used the ‘rda’ function of the R package ‘vegan’ to decompose the standardized correlation matrix into PC axes 1 and 2. We used the ‘vegan’ function ‘envfit’ (*50*) to treat the 7 environmental PC axes as responses in multiple regressions with the PC axes as explanatory variables (environmental variable ~ PC axis 1 + PC axis 2) to produce *P-* and R^2^ values used to assess the extent of explained variation.

We used the ‘ols’ function of the ‘statsmodels’ Python package to run multiple linear regression with the command ‘smf.ols(‘bestR2 ~ 1 + AA + protein_binding + DNA_binding + RNA_binding + in_disorder’, data = data, hasconst = True).fit()’ where bestR2 is the squared correlation (R^2^) resulting from the environmental association analysis, 1 the intercept, AA the repeated amino acid in the tract (as a categorical variable), ‘protein_binding’ an ordinal variable with values 0, 1, 2, or 3 depending on the extent the three different PPI prediction tools agreed (for instance, 3 if all tools agreed), and ‘DNA_binding’, ‘RNA_binding’, and ‘in_disorder’ were treated as binary (0/1) variables.

The luciferase experiment was performed 12 times per construct on one day. We used a Welch T-test to test if the differences in mean luciferase outputs was statistically significant.

## Author contributions

KSJ and MAB conceived the study. WBR performed the analyses with input from JBK, TR, OKT, SJ, MAB and KSJ. IMS and A.G. performed experimental work. WBR wrote the manuscript with input from KSJ, MAB, SJ, JBK, TR, IMS, AG and OKT.

## Data and materials availability

Our STR variant calling of protein coding STRs in 770 accessions are available as S. File 1 at https://figshare.com/s/8d08c53db83db3f223c7. Disorder prediction of Arabidopsis protein sequences can be retrieved from D2P2.pro. BED files of predicted IDRs and disordered interaction regions, as well as the exact TAIR10 peptide sequences used are available as S. File 8 at https://figshare.com/s/8d08c53db83db3f223c7. The gene expression results used in this study are available in the Supplementary Materials of (*9*). The environmental data used in this study is available in (*17*). Python scripts used to model, perform statistical analysis and to generate the figures are available at https://figshare.com/s/8d08c53db83db3f223c7.

## Supplementary Materials

**S. Figure 1.**
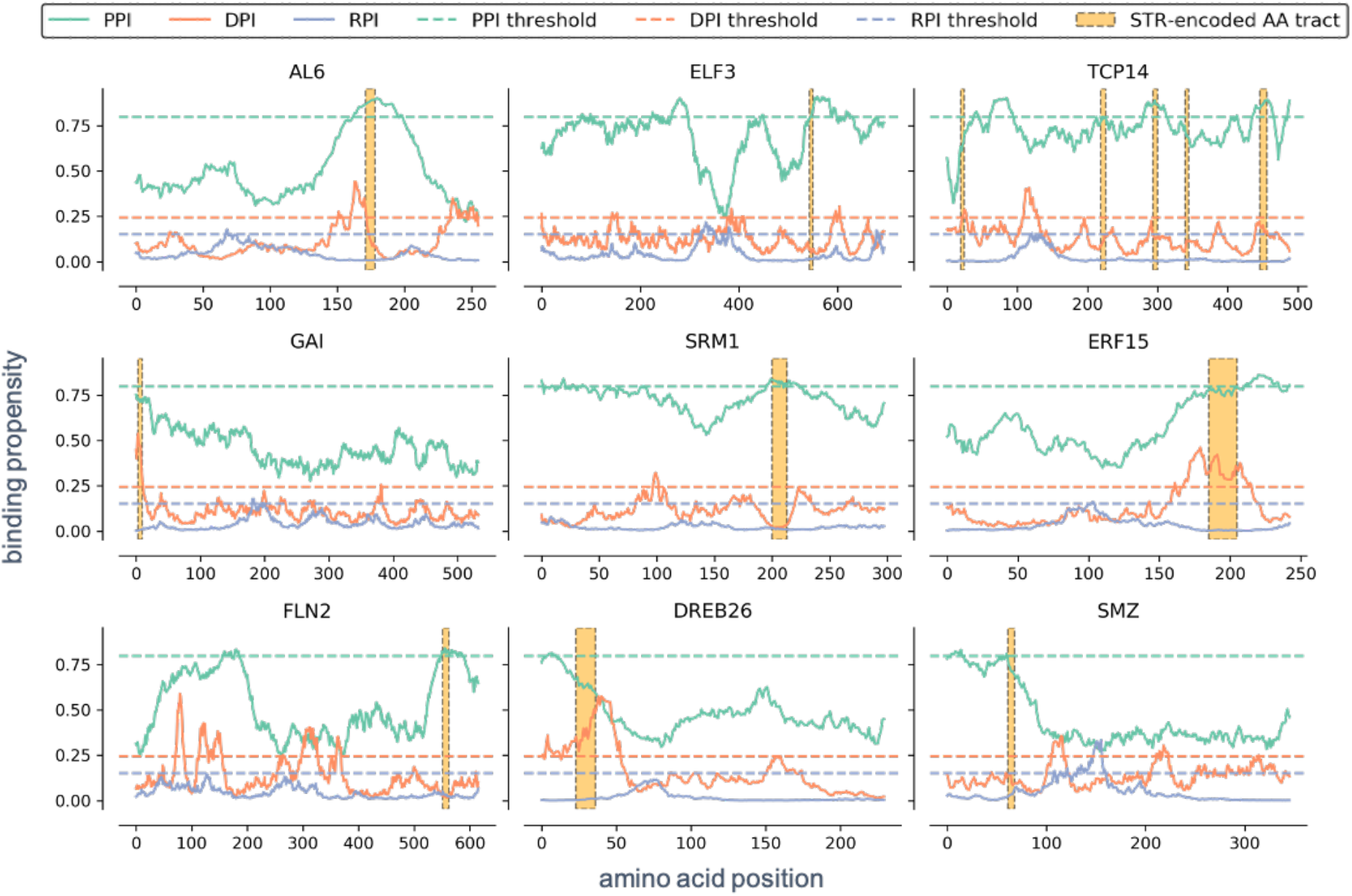
Examples of disordered interaction region prediction diagrams. The y-axis shows the binding propensity as predicted by DisoRDPbind. The x-axis shows the coordinates of the protein sequences. PPI: protein-protein interaction, DPI: DNA-protein interaction, RPI: RNA-protein interaction. Only predictions from DisoRDPbind are shown. The glutamate tract in ALFIN-LIKE 6 (AL6) influences the activation of its own promoter and the interaction with REALLY INTERESTING NEW GENE 1A (RING1A)^10^. Length variation in a glutamine tract in EARLY FLOWERING 3 (ELF3) correlates with thermal responsiveness^1^, and overlaps a predicted PPI region. TEOSINTE BRANCHED1, CYCLOIDEA, AND PROLIFERATING CELL FACTOR 14 (TCP14) contain many STR-encoded amino acid tracts, and multiple of these were linked to elevated protein-protein binding propensity. GIBBERELLIC ACID INSENSITIVE (GAI), SALT-RELATED MYB 1 (SRM1), ETHYLENERESPONSIVE TRANSCRIPTION FACTOR 15 (ERF15), FRUCTOKINASE-LIKE 2, CHLOROPLASTIC (FLN2), DEHYDRATION-RESPONSE ELEMENT-BINDING PROTEIN 26 (DREB26) and SCHLAFMUTZE all contain amino acid tracts that overlap elevated binding propensities.

**S. Figure 2.**
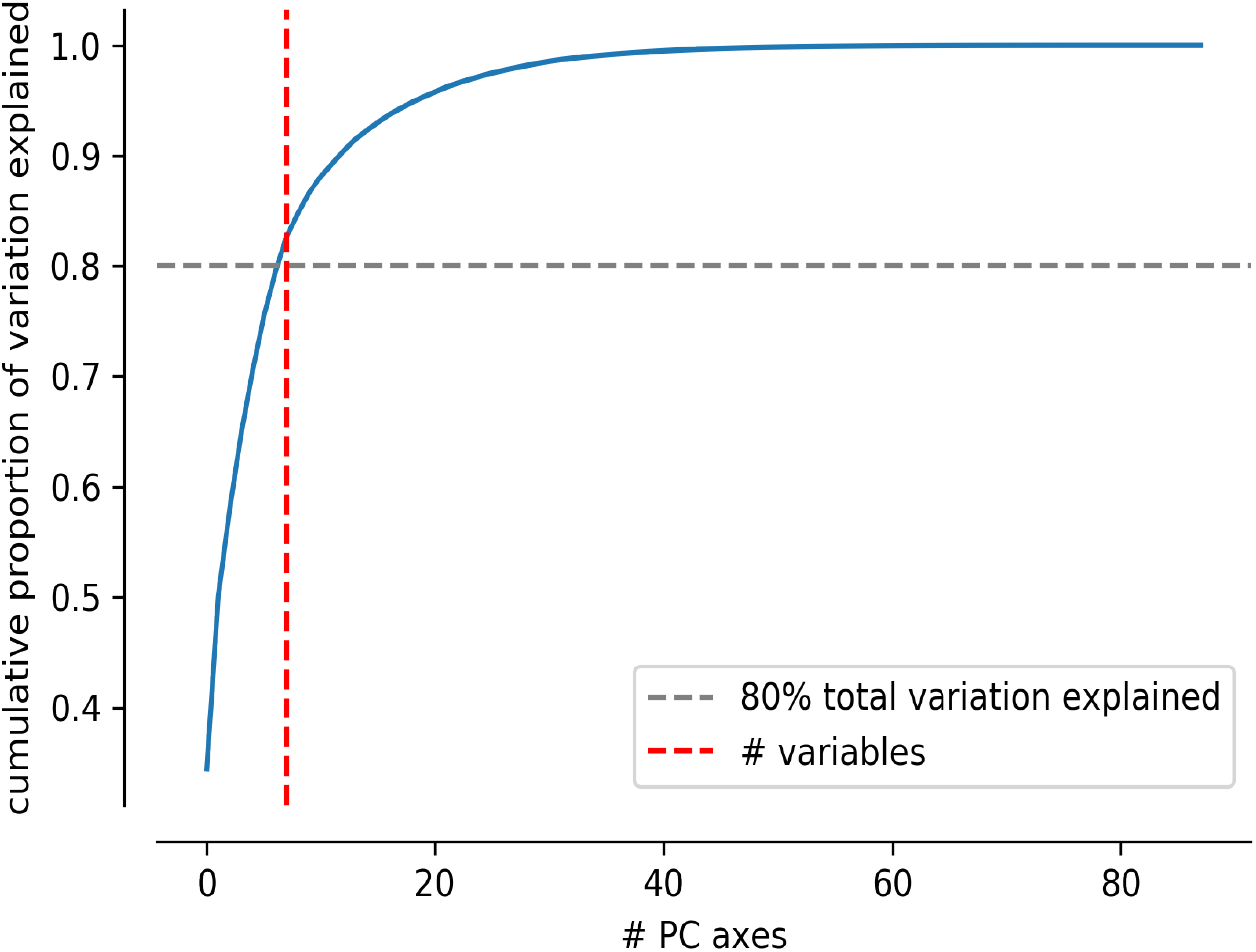
The cumulative proportion of environmental variation explained by principal component (PC) axes. The cumulative proportion of explained variation (y-axis) reached 0.8 when seven PC axes were included (x-axis: number of PC axes).

**S. Figure 3.**
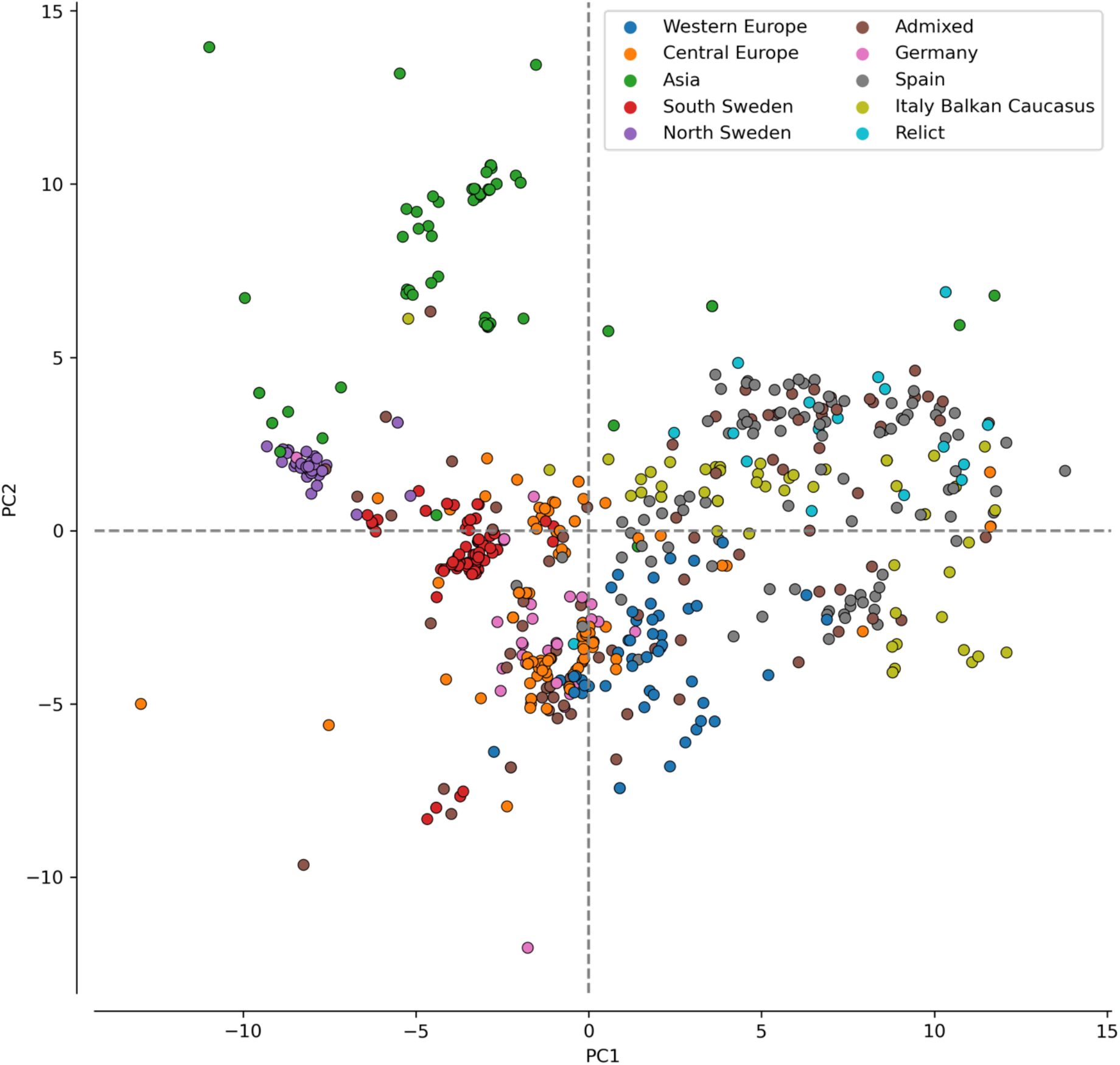
The position of accessions included in this study on principal component (PC) axis 1 and PC axis 2. The first axis captures 34.4% of the variation in the 88 environmental variables (listed in S. File 4), and the second axis captures 15.7% of the variation. Accessions are colored by the groups designated by the 1001 Genomes Consortium.

**S. Figure 4.**
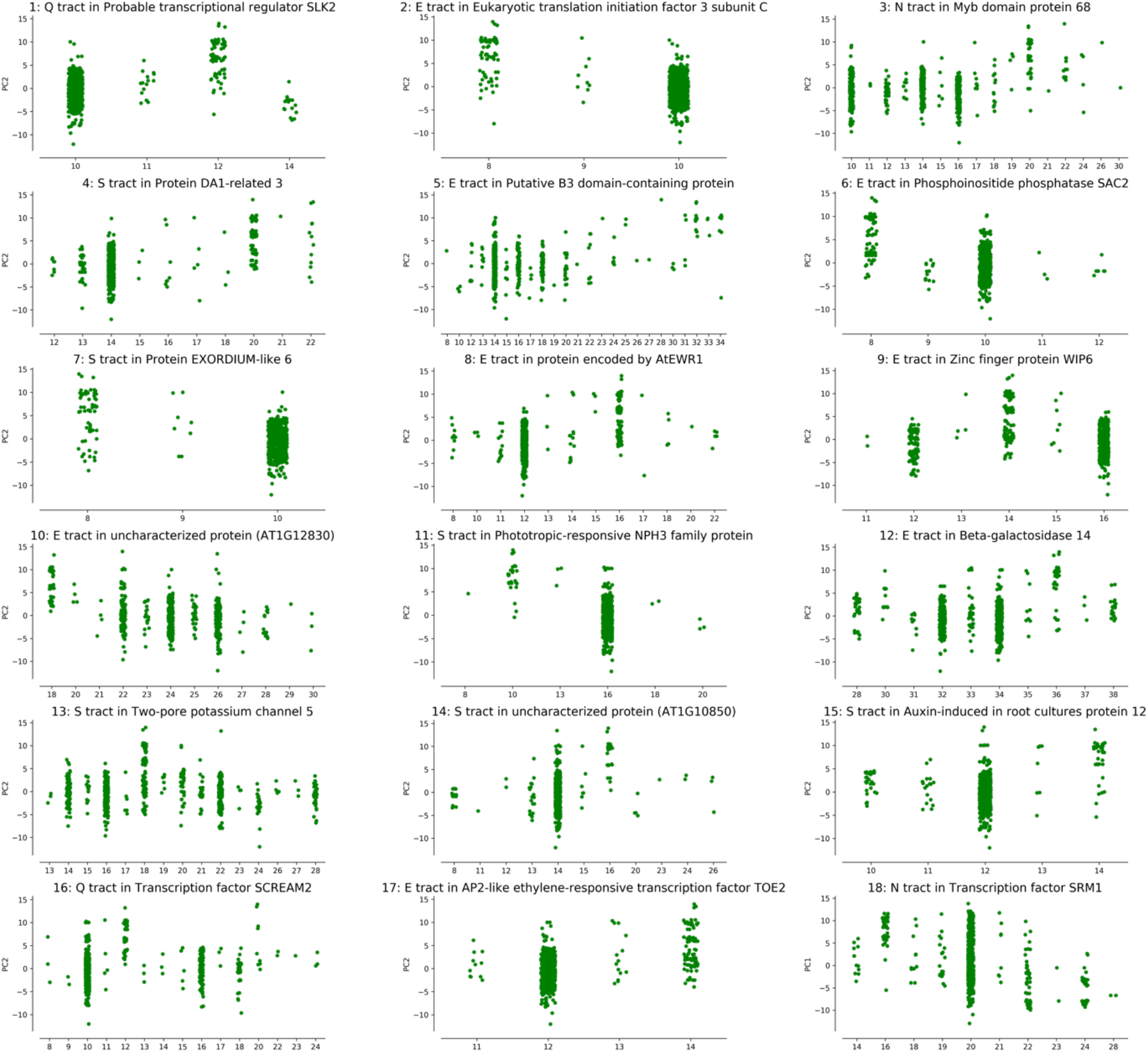
Top 18 associations (by *P-*value) between amino acid tract length variation in asparagine (N), serine (S), glutamate (E) and glutamine (Q) tracts and environmental variation. **(1)** Probable transcription regulator SEUSS-Like (SLK) 2 is involved in regulating abiotic stress response genes(*51*). Its poly-Q tract is in a predicted protein-protein interaction (PPI) region in a Q-rich region of the protein upstream of its dimerization domain. **(2)** Eukaryotic translation initiation factor 3 subunit C is a component of the eukaryotic translation initiation factor 3 complex. The poly-E tract is in an intrinsically disordered region (IDR) approximately in the middle of the protein sequence. **(3)** Myb domain protein 68 is directly involved in toleration of severe heat stress (*52*). The poly-N tract is located after a larger IDR in the C-terminal half of the protein. **(4)** Protein DA1-related 3 is a member of the DA1 gene family, involved in the control of seed and organ size (53). The poly-S tract is in the C-terminal half of the protein. **(5)** Putative B3 domain-containing protein is of unknown function. The poly-E tract is in a predicted PPI region located near the C-terminus, after the B3 DNA binding domain. **(6)** Phosphoinositide phosphatase suppressor of actin (SAC) 2 is a metabolic enzyme important for vacuolar morphology (54). The poly-S tract is in a predicted IDR located close to the C-terminus. **(7)** Protein EXORDIUM-like 6 is in the EXORDIUM family of genes, several of which are involved in promoting growth in response to brassinosteroids (*55*). The poly-S tract is in the signal peptide. **(8)** Enhancer of vascular Wilt Resistance 1 (EWR1) is a plant peptide that conveys resistance to *Verticillium dahliae,* a plant fungal pathogen (*56*). The poly-E tract is in an IDR located after the signal peptide. **(9)** Zinc Finger protein WIP6 (also known as DEFECTIVELY ORGANIZED TRIBUTARIES 5) is linked to leaf vein patterning(*57*). The poly-E tract is in a predicted PPI region downstream of a 4^th^ zinc finger, close to the C-terminus. **(10)** The protein encoded by AT1G12830 is completely disordered and has no known function. The poly-E tract is located in the middle of the protein. **(11)** Phototrophic-responsive NPH3 family protein does not have a known function. Its poly-S tract is in an IDR close to the N-terminus. **(12)** Beta-galactosidase 14 is a glycosidase involved in carbohydrate metabolism. The poly-E tract is in a predicted PPI region in its disordered, N-terminal tail. **(13)** Two-pore potassium channel 5 is a probable tonoplast ion channel(*58*). The poly-S tract is in a predicted PPI region near the C-terminus. **(14)** The protein encoded by AT1G10850 has a protein kinase domain, and a signal peptide, but has no known function. The poly-S tract is in a predicted DNA-protein interaction region in the beginning of the signal peptide. **(15)** Auxin-induced in root culture protein 12 is a plasma membrane cytochrome with a signal peptide(*59*). The poly-S tract is in the beginning of the signal peptide. **(16)** Transcription factor SCREAM 2 is a central regulator of stomatal development and is involved in cold acclimation (*60, 61*). Its poly-Q tract is in an IDR in the N-terminal half of the protein sequence. **(17)** AP2-like ethylene-responsive transcription factor TARGET OF EARLY ACTIVATION (TOE) 2 regulates flowering time and innate immunity (*62, 63*). The poly-E tract is in a predicted PPI in the N-terminal half of the protein. **(18)** Transcription factor SALT-RELATED MYB (SRM) 1 regulates the response to salt stress (*64*). The protein contains a poly-N tract in a predicted PPI in the C-terminal half of the protein.

**S. Figure 5.**
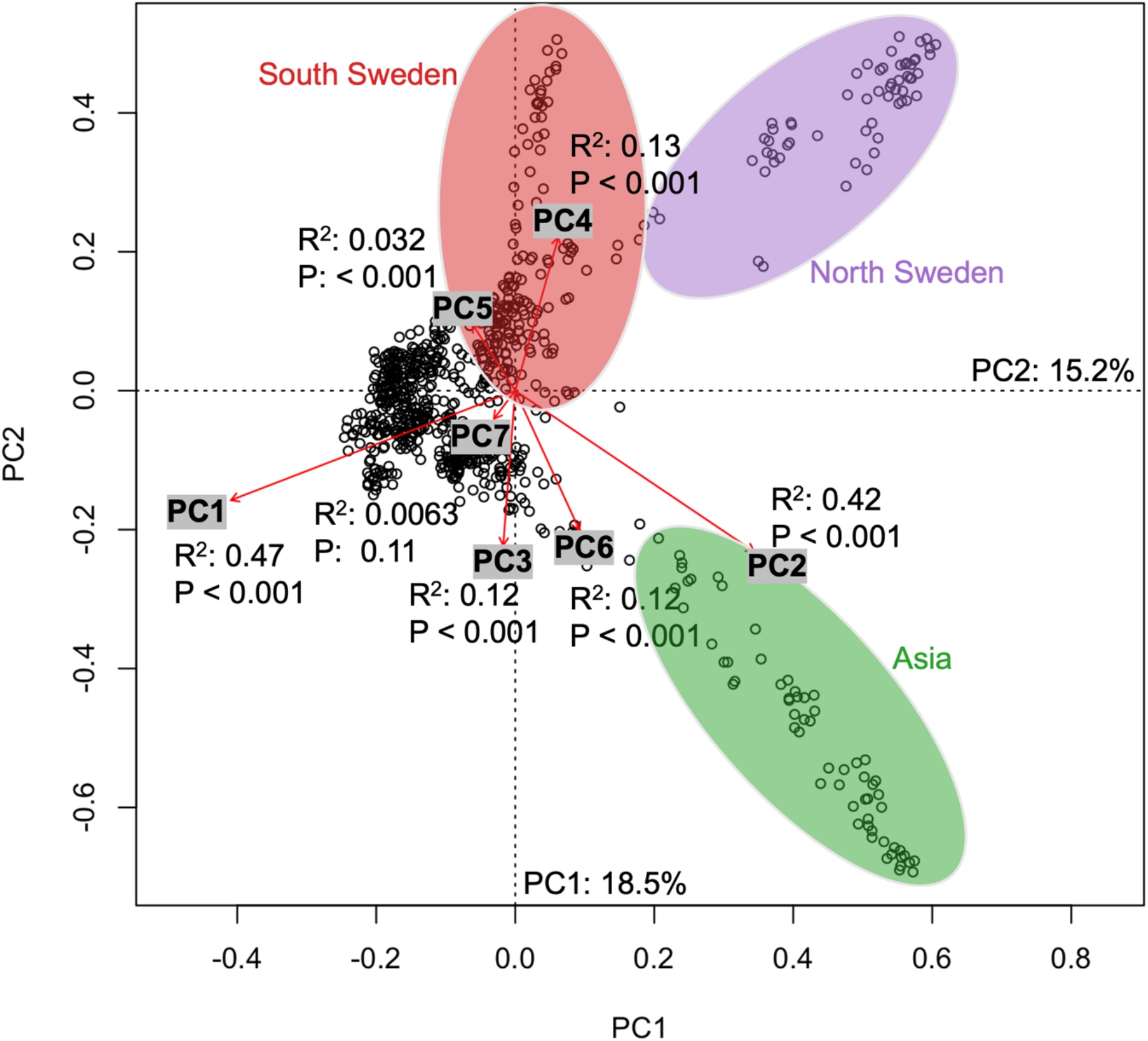
Putative neutral markers are significantly structured by environmental principal component (PC) axes 1 and 2. PC axis 1 (x-axis) and PC axis 2 (y-axis) represents gradients that best explain the structuring of accessions, based on putative neutral SNPs. Red arrows denote the environmental PC axes projected in the genetic space. Arrows are scaled with R^2^, and indicate to what extent the genetic structure can be explained by the environmental PC axes. For clarity, the three most prominent clusters (South Sweden, North Sweden and Asia) are highlighted (a few non-South Sweden accessions are present in the South Sweden cluster).

**S. Figure 6.**
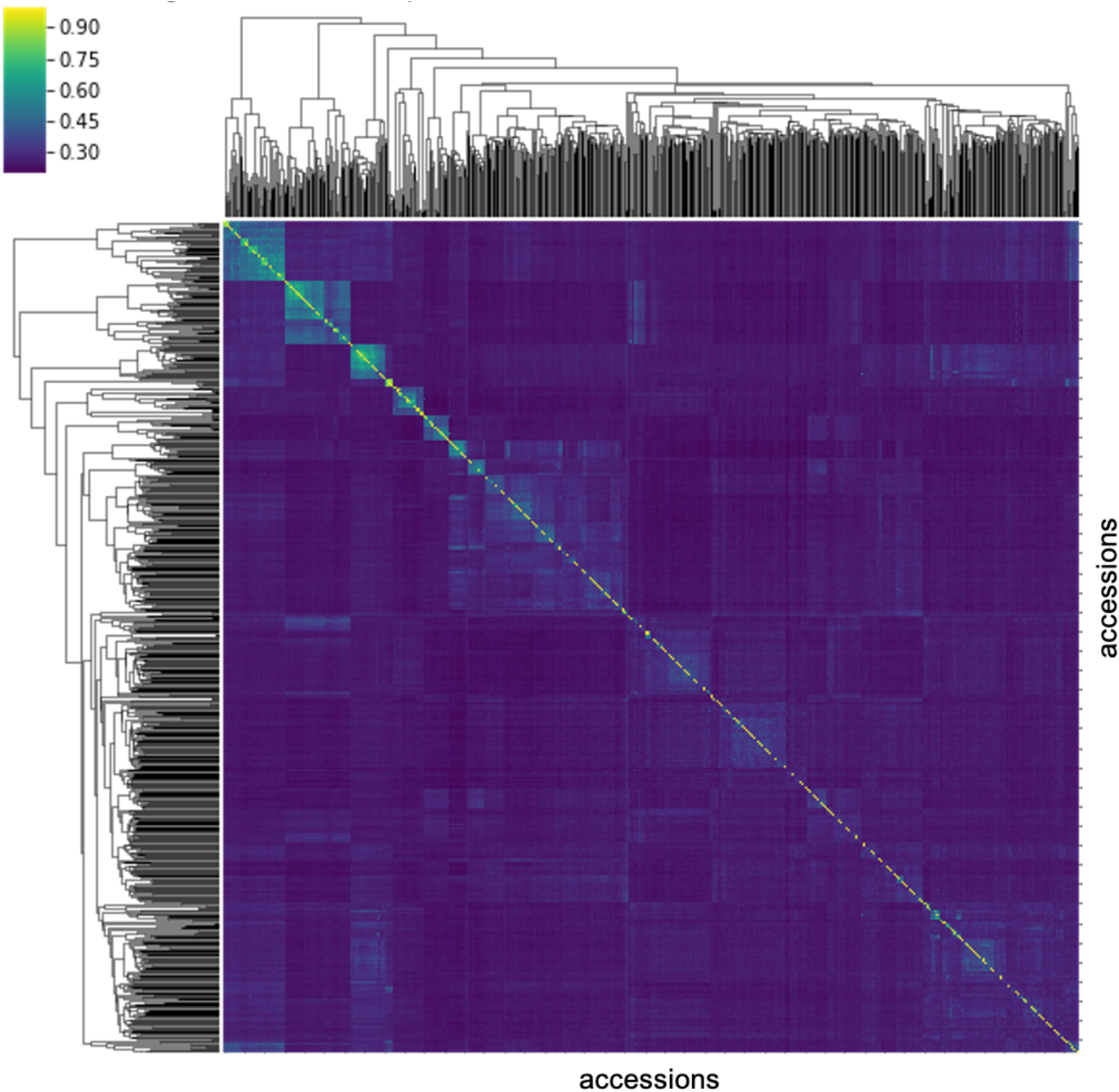
Standardized pairwise correlation matrix of putative neutral genetic markers. Higher values (from dark blue to yellow) denote more genetically similar pairs of accessions.

**S. Figure 8.**
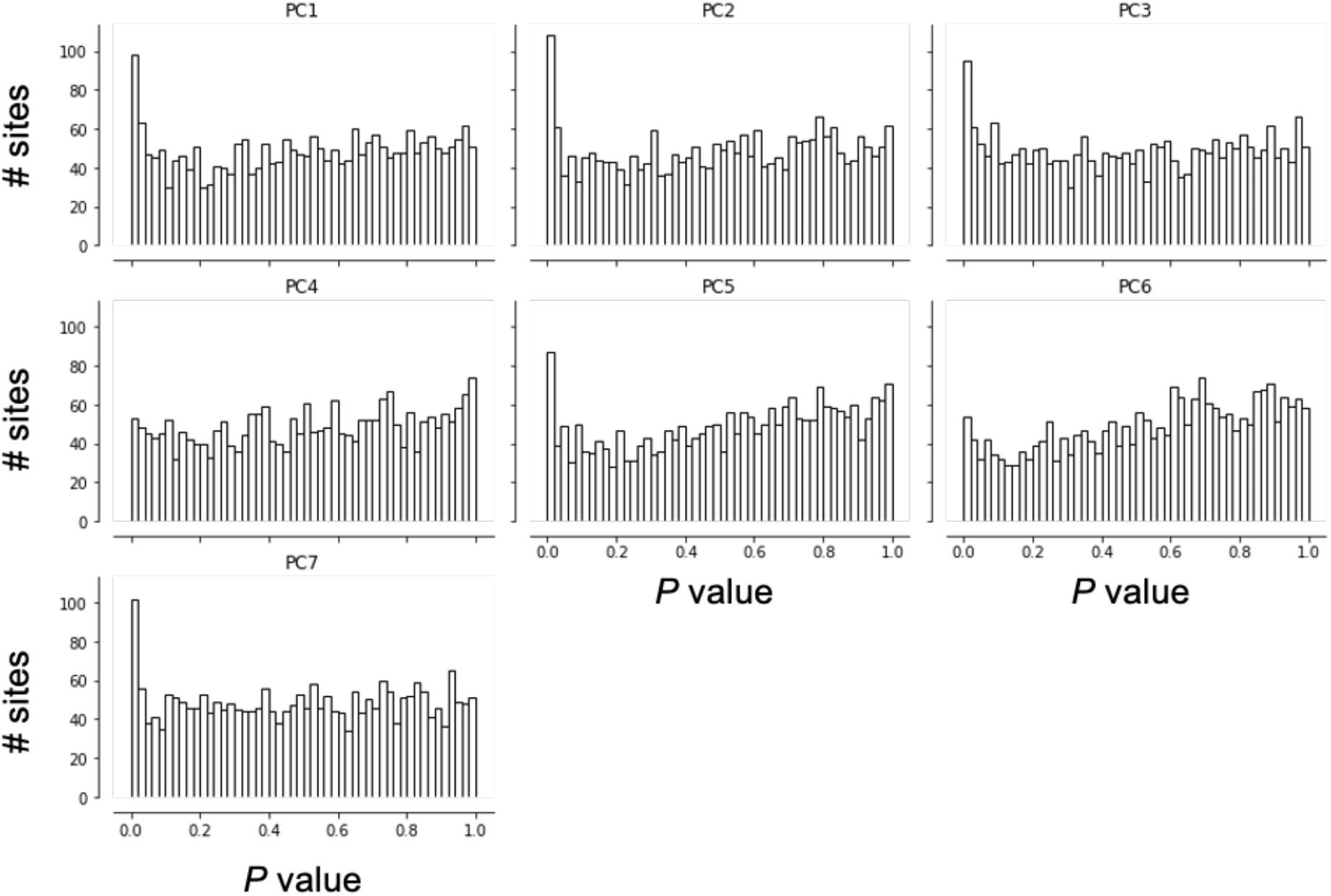
*P*-value histograms resulting from environmental association analyses with correction for putative neutral genetic markers.

**S. Table 1.** The number of different STR-encoded amino acid tracts in the *A. thaliana* proteome (source data for Figure 1A). Numbers were based on the primary transcript only (ATXXXXXXX.1), to avoid duplicate counts of tracts.

| repeated amino acid in tract | # tracts |
| --- | --- |
| A | 62 |
| C | 2 |
| D | 337 |
| E | 532 |
| F | 50 |
| G | 221 |
| H | 126 |
| I | 2 |
| K | 295 |
| L | 41 |
| M | 21 |
| N | 166 |
| P | 210 |
| Q | 174 |
| R | 21 |
| S | 499 |
| T | 82 |
| V | 11 |
| Y | 2 |

**S. Table 2. Source data and results of Fisher’s exact test of dependence between amino acids encoded by STRs and STRs in predicted disorder or predicted disordered binding sites.** Available at https://figshare.com/s/8d08c53db83db3f223c7

**S. Table 3. Results from running Gene Ontology and protein family (PFAM) enrichment analysis via the STRING database.** Available at https://figshare.com/s/8d08c53db83db3f223c7

**S. Table 4. Source data and results of Fisher’s exact test of dependence between STRs in predicted disorder or predicted disordered binding sites and being associated with differences in gene expression levels.**

Available at https://figshare.com/s/8d08c53db83db3f223c7

**S. Table 5.** Top negative and positive correlations between individual environmental variables and environmental principal component (PC) axis 1-7. A list of all variables used in the PC analysis are available as S. File 3. The variables were collected by Ferrero-Serrano and Assmann (2019) and further descriptions of the variables can be found in their study.

| correlation<br>(Pearson's R) | variable | category | PC<br>axis |
| --- | --- | --- | --- |
| -0.82 | spring frost days (CRU Frs spring) | temperature | 1 |
| 0.92 | spring maximum temperature (CRU Tmx spring) | temperature | 1 |
| -0.80 | net primary production (NPP based on GPCP) | productivity | 2 |
| 0.71 | temperature seasonality (CHELSA BIO4) | temperature | 2 |
| -0.55 | net primary production (dNPPdP) | productivity | 3 |
| 0.69 | spring precipitation (CRU Pre spring) | precipitation | 3 |
| -0.51 | mean diurnal temperature range (CHELSA BIO2) | temperature | 4 |
| 0.51 | mean annual precipitation (CRU precipitation) | precipitation | 4 |
| -0.33 | mean temperature, wettest quarter (CHELSA BIO8) | temperature | 5 |
| 0.82 | soil toxicity (GAEZ Toxicity) | edaphic | 5 |
| -0.28 | isothermality (CHELSA BIO3) | temperature | 6 |
| 0.71 | mean annual precipitation (GPCC Fulldata Precipitation) | precipitation | 6 |
| -0.49 | spring minimum temperatures (CHELSA Tmin spring) | temperature | 7 |
| 0.30 | soil carbon density (ISRIC WISE Soil Carbonate Density) | edaphic | 7 |

**S. Table 6. Accessions’ values on the environmental PC axes 1-7.** Available at https://figshare.com/s/8dO8c53db83db3f223c7

**S. Table 7. Source data and results of Fisher’s exact test of dependence between the type of amino acid and the association with environmental PC axes 1-7.** Available at https://figshare.com/s/8d08c53db83db3f223c7

**S. Table 8.**
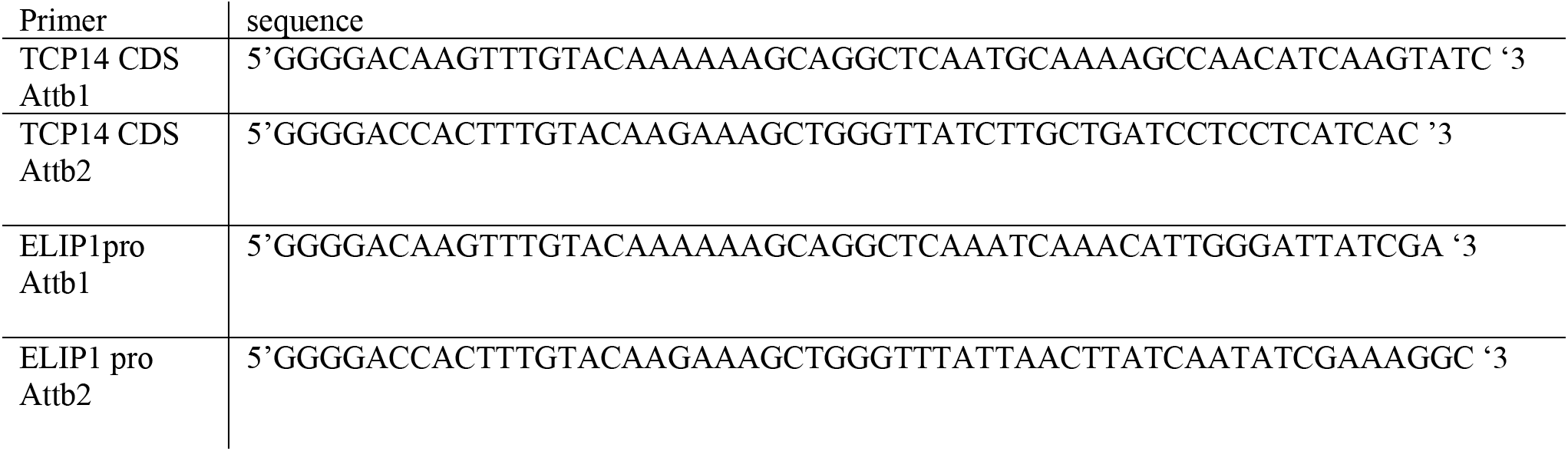
Primers used in this study.

**S. File 1. Coding STR genotype matrix of diploid short tandem repeat unit counts, 770 accessions.** Available at https://figshare.com/s/8d08c53db83db3f223c7

**S. File 2. STRING database network analysis of proteins with STR-encoded amino acid tracts in predicted IDRs or predicted disordered interaction regions.** Available at https://figshare.com/s/8d08c53db83db3f223c7

**S. File 3. Results of sequencing *TCP14* (Col-0) and *TCP14* (CS77239).** Available at https://figshare.com/s/8d08c53db83db3f223c7

**S. File 4. The 88 environmental variables with complete data on the 770 accessions analyzed in this study.** See Ferrero-Serrano and Assmann (2019) for further descriptions of the variables. Available at https://figshare.com/s/8d08c53db83db3f223c7

**S. File 5. Results of the environmental association analysis.** Available at https://figshare.com/s/8d08c53db83db3f223c7

**S. File 6. Results of the environmental association analysis, mock STR genotypes.** Available at https://figshare.com/s/8d08c53db83db3f223c7

**S. File 7. Results of the environmental association analysis with population structure correction.** Available at https://figshare.com/s/8d08c53db83db3f223c7

**S. File 8. IDR and disordered interaction region BED files and the TAIR10 peptide sequences used in this study.**

